# NOD2 modulates MDA5 signaling to promote coxsackievirus B3 replication

**DOI:** 10.1101/2025.05.06.652463

**Authors:** Sharon K. Kuss-Duerkop, Lauren C. Atencio, Elizabeth A. Nail, Ian L. Sparks, Thao N. Huynh, Brian C. Russo, A. Marijke Keestra-Gounder

## Abstract

Coxsackievirus is an enteric virus that encounters intestinal epithelial cells (IECs) after ingestion and can cause a range of illnesses from mild to severe, including gastroenteritis, pancreatitis, central nervous system diseases and myocarditis. Sensing of CVB3 RNA by melanoma differentiation-associated gene (MDA5) induces a Type 1 Interferon (T1IFN) response that is critical for reducing viral replication and dissemination within cells and the host. The cytosolic nucleotide oligomerization domain-containing protein 2 (NOD2), however, has been implicated in a proviral role in CVB3-induced myocarditis, but the underlying mechanism is not well understood. Because NOD2 is especially important in maintaining intestinal immune homeostasis and CVB3 is an enteric virus that initially infects the intestinal tract leading to virus dissemination to other tissues, we sought to determine whether NOD2 expressed in IECs would impact CVB3 infection. Here, we demonstrate that IEC NOD2 promotes CVB3 replication. Replication of a related *Enterovirus,* poliovirus, was also enhanced by NOD2. We discovered that *NOD2^−/−^* IECs have increased T1IFNs and interferon stimulated gene (ISG) expression, including *IFIH1*, the gene encoding for MDA5. Replication of CVB3 was rescued when MDA5 expression was reduced in *NOD2^−/−^* IECs. CVB3 replication was further increased, and the T1IFN response reduced, in *NOD2^+/+^*IECs treated with the mitophagy inducer carbonyl cyanide m-chlorphenyl hydrazone (CCCP). Mitophagy induction in *NOD2^−/−^* IECs, however, had no effect on CVB3 replication, or the T1IFN response, suggesting a role for NOD2 in mitophagy. Overall, our data demonstrate that NOD2 promotes CVB3 replication in IECs by suppressing T1IFN expression and MDA5 activation via NOD2-mediated mitophagy. Since T1IFN restricts enteroviruses *in vivo*, CVB3 and other enteroviruses might exploit NOD2 activation in the intestinal tract to evade antiviral T1IFN responses. This evasion could enhance their replication in the intestine and promote their spread to other organs, potentially leading to diseases in the pancreas, heart, and central nervous system.

**Importance:** Coxsackievirus B3 (CVB3) is a significant human intestinal virus capable of spreading to other organs. It is closely linked to the development of type I diabetes and can cause severe heart infections, particularly in children. Our research demonstrates that CVB3 exploits the immunity protein NOD2, which usually protects against infections, to infect and replicate in intestinal cells. NOD2 also enhances the replication of poliovirus, another related human enteric virus. When expressed in the intestinal epithelial cells, NOD2 reduces the immune response to certain viral infections by inhibiting expression of the viral sensor MDA5. Overall, CVB3 benefits from NOD2-mediated dampening of antiviral immunity to promote its replication in intestinal cells during fecal-oral transmission.

## Introduction

NOD2 is a pattern recognition receptor (PRR) that was first identified to recognize and respond to muramyl dipeptide (MDP) moieties within peptidoglycan of Gram-positive and Gram-negative bacteria ^1–4^. Since, NOD2 has been shown to respond to a variety of peptidoglycan-deficient pathogens and activation of cellular processes often usurped by many different pathogens ^5–7^. Stimulation of NOD2 triggers RIP2K activation and the assembly of the nodosome, an innate immune complex that initiates NFκB signaling ^6, 8^. Viruses use host proteins to productively infect cells and complete their replication cycles. While NOD2 has been shown to exhibit broad antiviral activity against respiratory syncytial virus (RSV) ^9^, vesicular stomatitis virus (VSV) ^9^, influenza A virus (IAV) ^9–11^, foot and mouth disease virus (FMDV) ^12^ and human cytomegalovirus (CMV) ^13, 14^, it can also facilitate infections by certain arboviruses, SARS-CoV-2, respiratory coxsackievirus B5 (CVB5) ^15^ and enteric coxsackievirus B3 (CVB3) ^16^.

CVB3 is a positive-sense, single-stranded RNA *Enterovirus* in the Family *Picornaviridae*, which causes a myriad of diseases including gastrointestinal (GI) sequelae, myocarditis, pancreatitis and central nervous system diseases, such as encephalitis ^17–20^. CVB3 infections may also trigger type I diabetes in susceptible individuals ^21, 22^. CVB3 infects via the fecal-oral route and can then disseminate to other organs where it replicates robustly and initiates disease ^19^. NOD2 exacerbates CVB3-induced myocarditis in mice, and *Nod2^−/−^* mice displayed decreased inflammation, apoptosis, CVB3 receptor expression and viral replication in cardiac tissues at day seven post-infection compared to wild-type mice when virus was administered intraperitoneally ^16^. Tschöpe *et al.* (2017) also demonstrated that *NOD2* expression, as well as *NLRP3*, was increased in hearts of myocarditis patients with confirmed CVB3 infection ^16^. This study indicated a pathogenic effect of NOD2 in the heart and underscored that cardiac NOD2 is a critical determinant of CVB3 disease severity. Similarly, *NOD2* expression was induced in human fetal brain cells infected with Zika virus (ZIKV). Chemical inhibition of NOD2 reduced replication of ZIKV, dengue virus (DENV), Mayaro virus (MAYV), SARS-CoV-2 and CVB5 by enhancing type I interferon (T1IFN) responses ^15^. Gene expression of interferon-stimulated genes (ISGs), including *NLRC4*, *NLRC5*, *OAS1*, *IFIT1* and *MX1*, was reduced by NOD2 inhibitors in lung epithelial cells ^15^. These two reports indicate that NOD2 promotes some viral infections by modulating immunity. Conversely, NOD2 inhibits other viral infections by enhancing mitochondrial antiviral signaling protein (MAVS)-dependent T1IFN responses to single-stranded RNA (ssRNA) as well as negative-sense RNA viruses in human embryonic kidney 293 cells, bone marrow-derived macrophages (BMDM), A549 lung epithelial cells, mouse embryonic fibroblasts (MEF) and normal human bronchial epithelial (NHBE) cells ^9^. *NOD2* expression was induced by RSV infection, and *Nod2*-deficient cells did not produce as much IFN-β as wild-type cells during RSV infection ^9^. *Nod2^−/−^* mice are more protected from RSV ^9^ and IAV infections ^10, 11^. Altogether, these data present a paradox for the role of NOD2 during viral infections. Understanding how NOD2 modulates innate immunity to diverse viruses could explain the contradictory outcomes to various viral infections and whether NOD2 would play a protective or pathogenic role.

T1IFN, which is commonly elicited after viral RNA sensing by the cytosolic RIG-I-like receptors (RLR): melanoma differentiation-associated gene (MDA5), retinoic acid-inducible gene-I (RIG-I) and laboratory of genetics and physiology 2 (LGP2), is critical for reducing viral replication and dissemination within cells and hosts ^23–25^. Picornaviruses, like CVB3, are sensed by MDA5, which is critical for antiviral T1IFN responses to CVB3 infection ^26–29^. Whereas it is generally accepted that MDA5 recognizes and responds to long double-stranded RNA (dsRNA), which is generated during replication of picornaviruses ^30–33^, other studies suggest more specific MDA5 ligands based on ssRNA sensing ^34–36^ or the combination of ssRNA and dsRNA ^37^. CVB3 viral RNA lacks the canonical RIG-I ligand 5’-di- or -triphosphate RNA commonly generated during some viral infections ^38, 39^, but recent evidence demonstrates that RIG-I also influences CVB3 infections ^40^. Moreover, RIG-I is cleaved during infections with picornaviruses, including CVB3 ^41,42^, suggesting that RIG-I impacts picornavirus infections by an unidentified mechanism. LGP2 acts to modulate RIG-I and MDA5 responses to virus infections, but it lacks a CARD domain and cannot initiate downstream signaling on its own ^25^. Viral RNA recognition by RLRs on the carboxy terminal end initiates conformational changes in MDA5 or RIG-I that expose the two amino terminal CARD domains, which interact with the MAVS CARD domain. This interaction with mitochondrial-associated MAVS results in downstream activation of both IRF3 and NFκB, which are transcription factors that translocate to the nucleus and elicit expression of T1IFNs, interferon α and β (IFNαβ). IFNαβ bind to the interferon αβ receptor (IFNAR) and activate JAK/STAT signaling, which induces expression of ISGs to establish an antiviral cellular state ^24^. CVB3 is sensitive to T1IFNs ^26, 43–46^ and encodes mechanisms to cleave important antiviral signaling proteins that diminish T1IFN signaling ^42, 47^. To prevent T1IFN-dependent tissue damage often observed in autoimmune diseases, the MDA5-MAVS signaling pathway is rapidly downregulated shortly after viral infection. This downregulation occurs through the interaction of MAVS with the E3 ubiquitin ligase RNF34 on the mitochondria, leading to MAVS polyubiquitination. The ubiquitin chains on MAVS are detected by the cargo receptor NDP52, which facilitates the recruitment of damaged mitochondria containing MAVS aggregates to the vacuole for degradation via autophagy. This process establishes a negative feedback loop that inhibits MAVS-mediated T1IFN signaling ^48^. Notably, the NOD2/RIPK2 signaling pathway regulates mitophagy by phosphorylating the mitophagy inducer ULK1 ^10^, and CVB3-induced mitophagy inhibits the T1IFN response by abrogating the MAVS/TBK1 interaction ^49^. However, the impact of NOD2 on T1IFN production and antiviral activity remains unclear.

Here, we explored the role of intestinal NOD2 during CVB3 infection. CVB3 initially infects IECs in the GI tract ^19^, where NOD2 plays a critical role in intestinal immunity ^50^. Consistent with findings of NOD2 involvement in supporting CVB3 replication during heart infections ^16^, we observed that intestinal NOD2 similarly facilitates CVB3 replication. The absence of NOD2 in IECs led to increased T1IFN gene expression, which restricted CVB3 replication. Moreover, CVB3 replication was restored in *NOD2^−/−^* cells when *IFIH1* (MDA5), but not *DDX58* (RIG-I), expression was silenced using small interfering RNAs (siRNAs). Induction of mitophagy suppressed the T1IFN response and enhanced viral replication in wild-type cells, whereas mitophagy induction in *NOD2^−/−^* cells had no effect on ISG expression or viral replication. Together, these findings reveal a novel pathogenic role of IEC-expressed NOD2 in inhibiting MDA5-mediated antiviral responses.

## Results

### NOD2 is required for efficient CVB3 replication

To ascertain whether NOD2 affected CVB3 replication in IECs, we generated *NOD2^−/−^* HCT116 human colonic epithelial cells by use of CRISPR/Cas9 targeting of *NOD2* or no template control (NTC). To determine if *NOD2* was inactivated by CRISPR/Cas9 targeting, *NOD2^+/+^* (HCT116 wild-type cells infected with NTC gRNA lentivirus) and *NOD2^−/−^* cells were stimulated with low and high concentrations of the NOD2 ligand MDP or the NOD1 ligand C12-iE-DAP. Quantitative RT-PCR (qRT-PCR) was performed for *IL8*, a pro-inflammatory gene stimulated by NOD2 signaling. Indeed, *NOD2^−/−^* cells did not produce *IL8* gene expression in response to MDP, whereas *NOD2^+/+^*cells displayed a strong response to MDP (Fig. S1A). Moreover, C12-iE-DAP stimulated NOD1 in *NOD2^−/−^* cells indicating that *NOD2^−/−^* cells could be stimulated by other PRR ligands and that the defect in *IL8* expression was specific to NOD2 stimulation (Fig. S1A). To test that loss of *NOD2* did not influence cell health, *NOD2^+/+^* and *NOD2^−/−^* cells were plated at the same density and cell growth was quantified out to 72 hours post-plating. We observed no difference in proliferation between *NOD2^+/+^* and *NOD2^−/−^* cells (Fig. S1B), and both cell lines could be cultured simultaneously for up to 30 passages with no apparent morphological or growth differences (data not shown). Indel analysis was performed to identify *NOD2* guide RNA g(RNA)/CAS9-induced mutations in *NOD2^−/−^* cells. Indeed, we identified an insertion mutation within the gRNA-targeted sequence of *NOD2*, near the end of the CARD2 domain, that would cause a frameshift and likely render all downstream domains non-functional. Additionally, we identified a mutation downstream of the gRNA sequence that could also affect the function of NOD2 (Fig. S1C). Next, *NOD2^+/+^* and *NOD2^−/−^* cells were infected with CVB3 for 24 or 48 hours, and CVB3 was quantified from either cells (cell-associated virus) or supernatants (extracellular virus). NOD2 enhanced the replication of CVB3 (Fig. 1A and Fig. S2A-C). To confirm that *NOD2^−/−^* cells were not altered in an off-target manner that affected CVB3 replication, we used GSK583, an inhibitor of the NOD2 signaling adapter RIPK2. GSK583 treatment of infected cells revealed that NOD2 signaling is critical for full replication of CVB3 in IECs (Fig. 1B and Fig. S2D). To ensure that NOD2 promotion of CVB3 replication was not specific to only HCT116 IECs, we examined CVB3 replication in other IECs treated with the RIPK2 inhibitor GSK583 to block NOD2 signaling. CVB3 replication was reduced when NOD2 signaling was inhibited in Caco-2 human colon epithelial cells (Fig. S3A) and MODE-K mouse small intestinal epithelial cells (Fig. S3B). Furthermore, mouse embryonic fibroblasts (MEFs) expressing NOD2 supported more CVB3 replication than MEFs lacking NOD2 (Fig. S3C). For all IECs, we confirmed that GSK583 concentrations used were non-toxic and would not inadvertently alter CVB3 replication by modulating cell health (Fig. S3D-F). Altogether, we generated a specific *NOD2^−/−^* HCT116 cell line that is more resistant to CVB3 infection and confirmed NOD2 specificity for the promotion of CVB3 infection.

**Figure 1.**
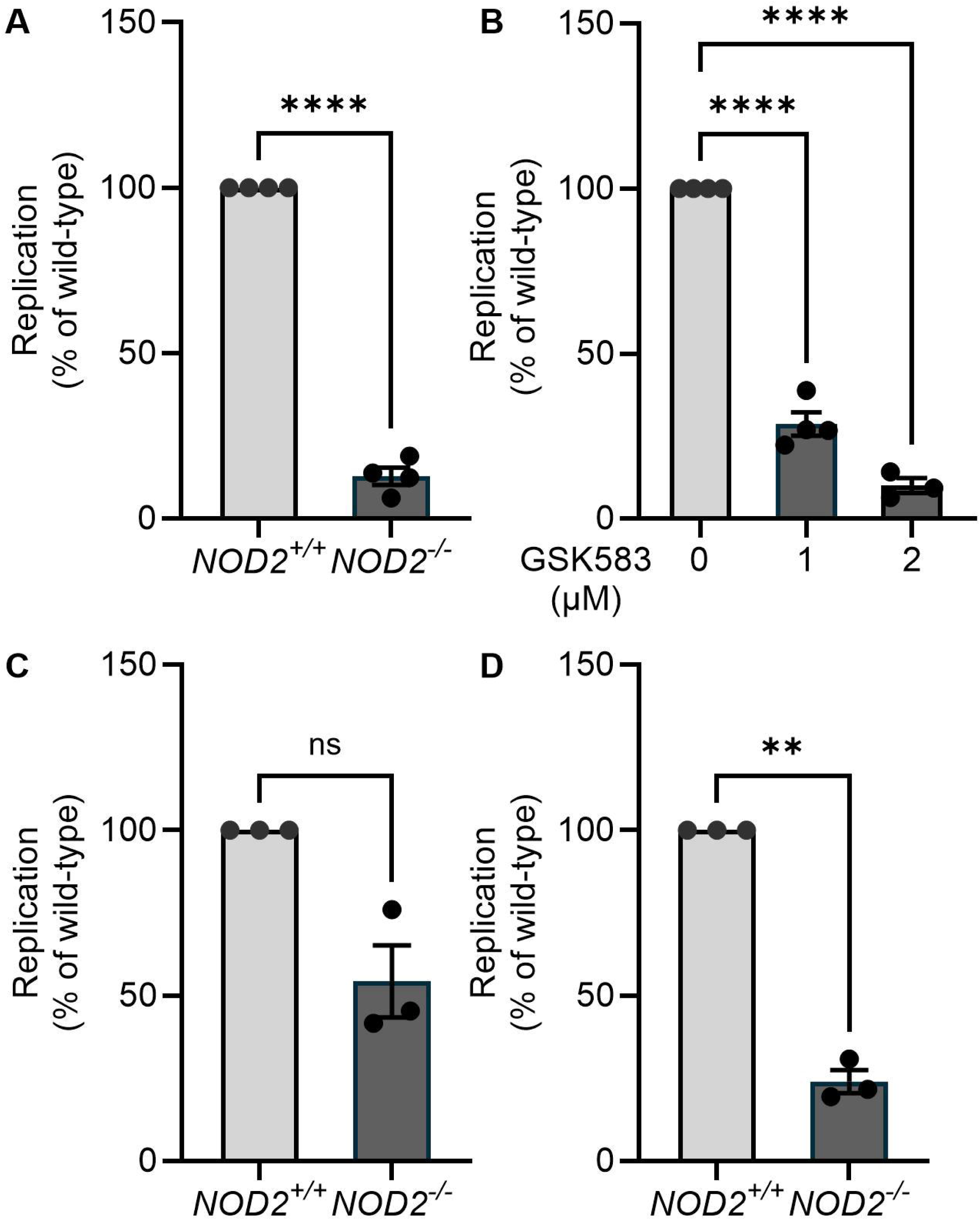
NOD2 promotes CVB3 replication in intestinal epithelial cells. (A) *NOD2^+/+^* or *NOD2^−/−^* HCT116 cells were infected with CVB3-H3 at an MOI=0.5 and cell-associated virus was quantified by plaque assay (N=4, duplicates). (B) HCT116 cells were pre-treated with 0.1% DMSO or the indicated concentrations of GSK583 for 20 minutes, and then infected at an MOI=0.5 in the presence of DMSO or GSK583 for 24 hours. At 24 hpi, cell-associated virus was quantified by plaque assay (N=3-4, duplicates). (C, D) *NOD2^+/+^* or *NOD2^−/−^* HCT116 cells were infected with CVB3-H3 for a single cycle of replication at (C) MOI=2 (N=3, duplicates) or (D) MOI=0.05 (N=3, duplicates). At 7 hpi, cell-associated virus was quantified by plaque assay. Data are displayed as percent virus compared to the wild-type control. \*\**p*<0.01, \*\*\*\**p*<0.0001, ns=not significant, unpaired *t*-test.

To help understand what mechanisms of resistance are present in *NOD2^−/−^* cells to reduce CVB3 replication, we performed a multiplicity of infection (MOI) test experiment using a single cycle of replication. We reasoned that if a restrictive host factor was upregulated in *NOD2^−/−^* cells to reduce CVB3 replication, then using a higher MOI might be able to rescue CVB3 replication by overwhelming or inhibiting the restrictive host factor. When *NOD2^+/+^* and *NOD2^−/−^*cells were infected for a single-cycle of replication at a high MOI (2), CVB3 replication was partially rescued in *NOD2^−/−^* cells (Fig. 1C and Fig. S2E), but not when a 40-fold lower MOI (0.05) was used (Fig. 1D and Fig. S2F). Therefore, NOD2 may restrict the function of a host factor that limits CVB3 replication. CVB3 replication during the high MOI infection was still trending lower in *NOD2^−/−^* cells compared to *NOD2^+/+^* cells (Fig. 1C), which suggests that the host factor was still partially active or other promotional roles of NOD2 exist to enhance CVB3 replication.

### NOD2 enhancement of CVB3 replication is independent of the viral receptor

A previous report demonstrated that expression of the CVB3 receptor CAR (gene: *CXADR*) was reduced in *Nod2^−/−^* compared to *Nod2^+/+^* mouse cardiac tissue, which could explain the reduction of CVB3 loads in mouse hearts ^16^. To address whether CAR expression was altered in *NOD2^−/−^* IECs accounting for reduced CVB3 replication, we performed multiple experiments. First, we employed another medically important picornavirus (genus *Enterovirus*) that uses a different receptor. Poliovirus uses CD155, or the poliovirus receptor (PVR), for cell entry. NOD2 and RIPK2 promoted poliovirus replication as well (Fig. 2A and B). Therefore, NOD2 may be important for the replication of other enteroviruses. Interestingly, NOD2 restricts the replication of foot-and-mouth-disease virus (FMDV), which is a non-human picornavirus in a different genus, *Aphthovirus* ^12^. Exploring how NOD2 positively influences some viral infections while negatively influencing others is crucial for understanding the contribution of NOD2 in various viral infections.

**Figure 2.**
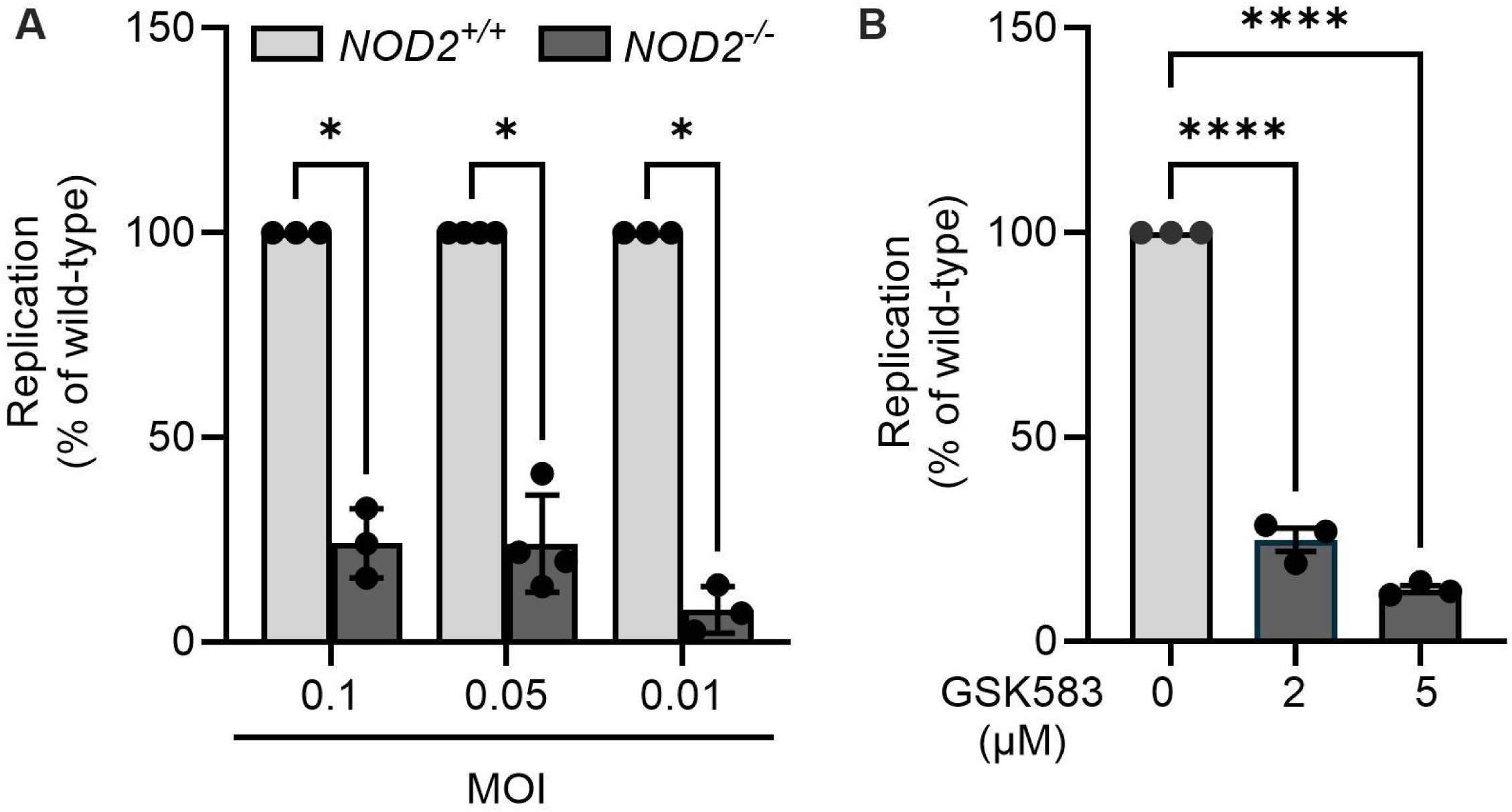
NOD2 promotes poliovirus replication in intestinal epithelial cells. (A) *NOD2^+/+^* or *NOD2^−/−^* HCT116 cells were infected with poliovirus at the indicated MOIs. 24 hpi, cell-associated poliovirus was quantified by plaque assay (N=3-4, duplicates). (B) HCT116 cells were pre-treated with 0.1% DMSO or the indicated concentrations of GSK583 for 20 minutes, and then infected with poliovirus in the presence of DMSO or GSK583 for 24 hours. At 24 hpi, cell-associated virus was quantified by plaque assay and displayed as percent virus compared to the wild-type control for each MOI (N=3). \**p*<0.05, \*\*\*\**p*<0.0001, unpaired *t*-test (A), One-Way ANOVA (B).

Because we observed that poliovirus replication was also reduced in *NOD2^−/−^* cells similar to CVB3 (Fig. 2), we presumed that reduction of CVB3 replication in *NOD2^−/−^* cells was not a result of altered CAR expression. To test this hypothesis, we examined *CXADR* gene expression and CAR protein levels by immunoblot. We did not observe transcript or protein level differences in *CXADR*/CAR expression between *NOD2^+/+^*and *NOD2^−/−^* cells (Fig. 3A and B), indicating that NOD2 enhances CVB3 replication at a step post-receptor binding. Expression of CAR may not differ between *NOD2^+/+^*and *NOD2^−/−^* cells, but perhaps cell surface expression of CAR is altered in *NOD2^−/−^* cells. Thus, we examined this using immunofluorescent staining of CAR on *NOD2^+/+^* and *NOD2^−/−^* cells and quantified the fluorescent signal per cell. No difference in plasma membrane expression of CAR was observed between *NOD2^+/+^* and *NOD2^−/−^* cells (Fig. 3C). Furthermore, to examine whether CVB3 receptor binding might be insufficient in *NOD2^−/−^* cells, we performed a CVB3 binding assay comparing *NOD2^+/+^* and *NOD2^−/−^* cells. The binding experiment with *NOD2^+/+^*and *NOD2^−/−^* cells was performed at either 37°C or 6°C under the premise that pre-binding CVB3 to cells at 6°C, during which internalization does not occur, would allow prolonged and sufficient receptor binding time in both cell lines, and therefore, might rescue the CVB3 replication defect in *NOD2^−/−^* cells. Room temperature virus was added to 37°C cells or cold virus was added to pre-chilled cells on ice. 37°C cells were incubated for 30 minutes at 37°C, and 6°C cells were incubated for one hour at 6°C. Following incubations, cells were washed twice and warm media was added for the duration of the 24-hour experiment. To confirm that CVB3 entry was reliant on CAR, cells were pre-treated and infected in the presence of isotype control or CAR antibodies. Indeed, blocking CAR with specific antibodies precluded CVB3 binding and entry (Fig. 3D), confirming that CAR is required for CVB3 infection. Our results indicate that pre-binding of CVB3 to *NOD2^+/+^* and *NOD2^−/−^* cells at 6°C did not increase CVB3 replication in *NOD2^−/−^* cells (Fig. 3D), suggesting that the CVB3 replication defect in *NOD2^−/−^* cells is mediated downstream of and independently of receptor binding.

**Figure 3.**
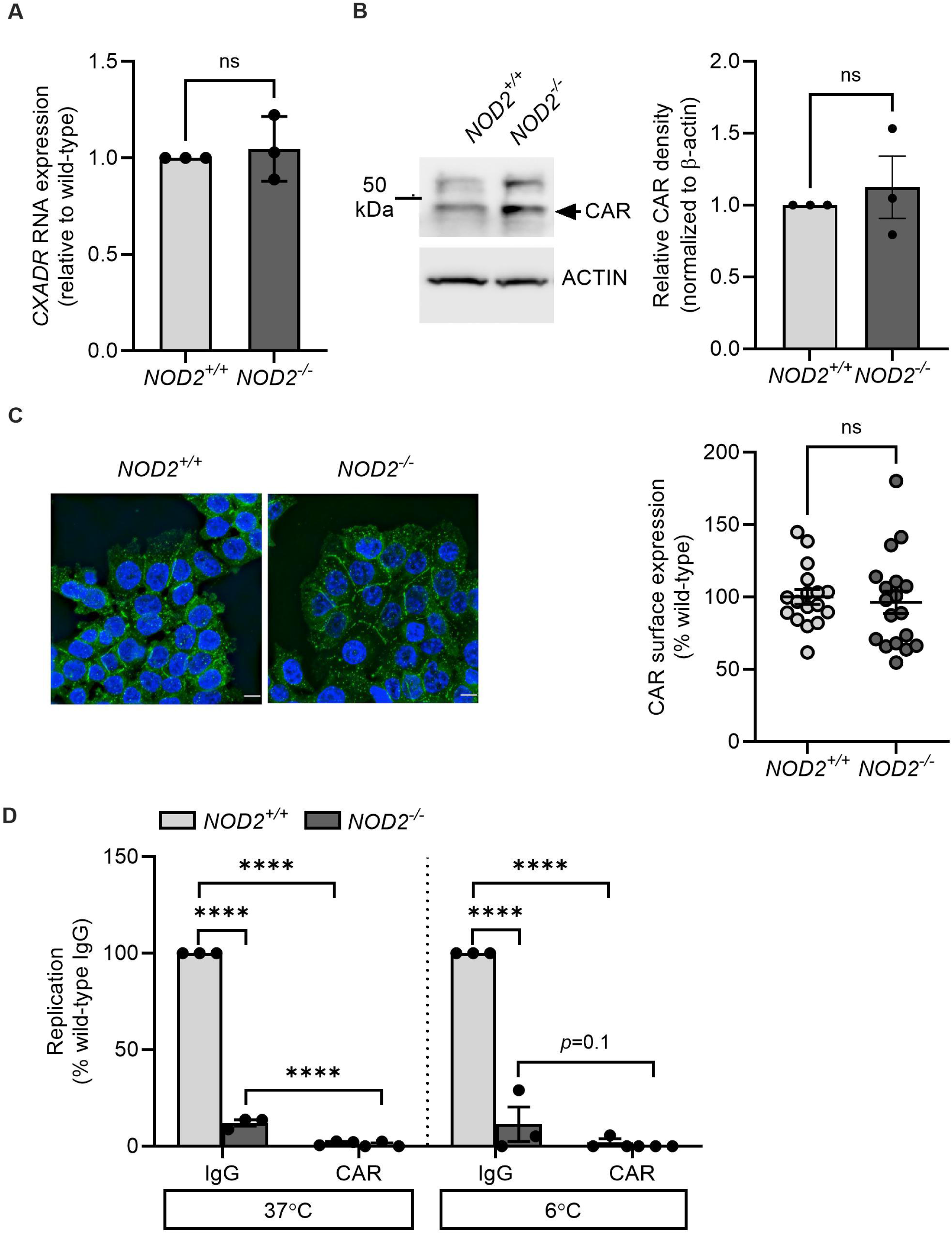
NOD2 does not affect CAR expression or CVB3 interactions with CAR. *NOD2^+/+^* or *NOD2^−/−^* HCT116 cells were plated and then collected 24 hours later. (A) qRT-PCR was performed for *CXADR* (N=3, duplicates) (paired *t*-test), (B) immunoblots were performed to determine CAR protein levels. Bar graph showing the relative protein density after normalization with β-actin (N=3, one representative image is shown) ns = not significant, paired *t*-test. (C) immunofluorescence assays were performed for surface-expressed CAR (N=3-4, representative images shown, scale bar = 10 μm) and quantified (N=3-4, 10-18 images each with 30-200 cells/frame) (unpaired *t*-test). (D) *NOD2^+/+^* or *NOD2^−/−^* HCT116 cells were plated and CVB3-H3 binding assays were performed as described in materials and methods (N=3, duplicates, \*\*\*\**p*<0.0001, two-way ANOVA).

### NOD2 suppresses type I interferon gene expression

Several lines of evidence indicate that NOD2 may regulate innate immunity during CVB3 infection ^15, 16^. Limonta *et al.* (2021) demonstrated that expression of several innate immune genes, including ISGs, increased in A549 lung epithelial cells when treated with the RIPK2 inhibitor GSK583 ^15^. We hypothesized that T1IFN gene expression may be increased in *NOD2^−/−^* cells compared to *NOD2^+/+^*cells, which could inhibit CVB3 and poliovirus replication. To investigate whether T1IFN gene expression was modulated by NOD2, *NOD2^+/+^* and *NOD2^−/−^* cells were plated at the same density, and expression of *IFNB*, *ISG15*, *ISG54, IFIH1* (MDA5), *DDX58* (RIG-I), and *IL8* was analyzed by qRT-PCR. T1IFN stimulated gene expression was significantly higher in *NOD2^−/−^* cells compared to *NOD2^+/+^* cells (Fig. 4A), indicating that baseline ISG expression is increased in *NOD2^−/−^* cells. NOD2 alteration of ISG expression was specific to T1IFN genes because expression of the pro-inflammatory chemokine *IL8* was unchanged in *NOD2^−/−^* cells compared to *NOD2^+/+^* cells. Intriguingly, protein expression of the viral RNA sensors MDA5 and RIG-I, encoded by *IFIH1* and *DDX58*, respectively, was significantly increased in *NOD2^−/−^*cells (Fig. 4A and S4B). Additionally, secreted IFNβ in the culture supernatant of unstimulated HCT116 cells was higher in *NOD2^−/−^* cells compared to *NOD2^+/+^*cells (Fig. 4B). Altogether, this suggests that *NOD2^−/−^* cells have increased antiviral responses that could contribute to reduced viral replication.

**Figure 4.**
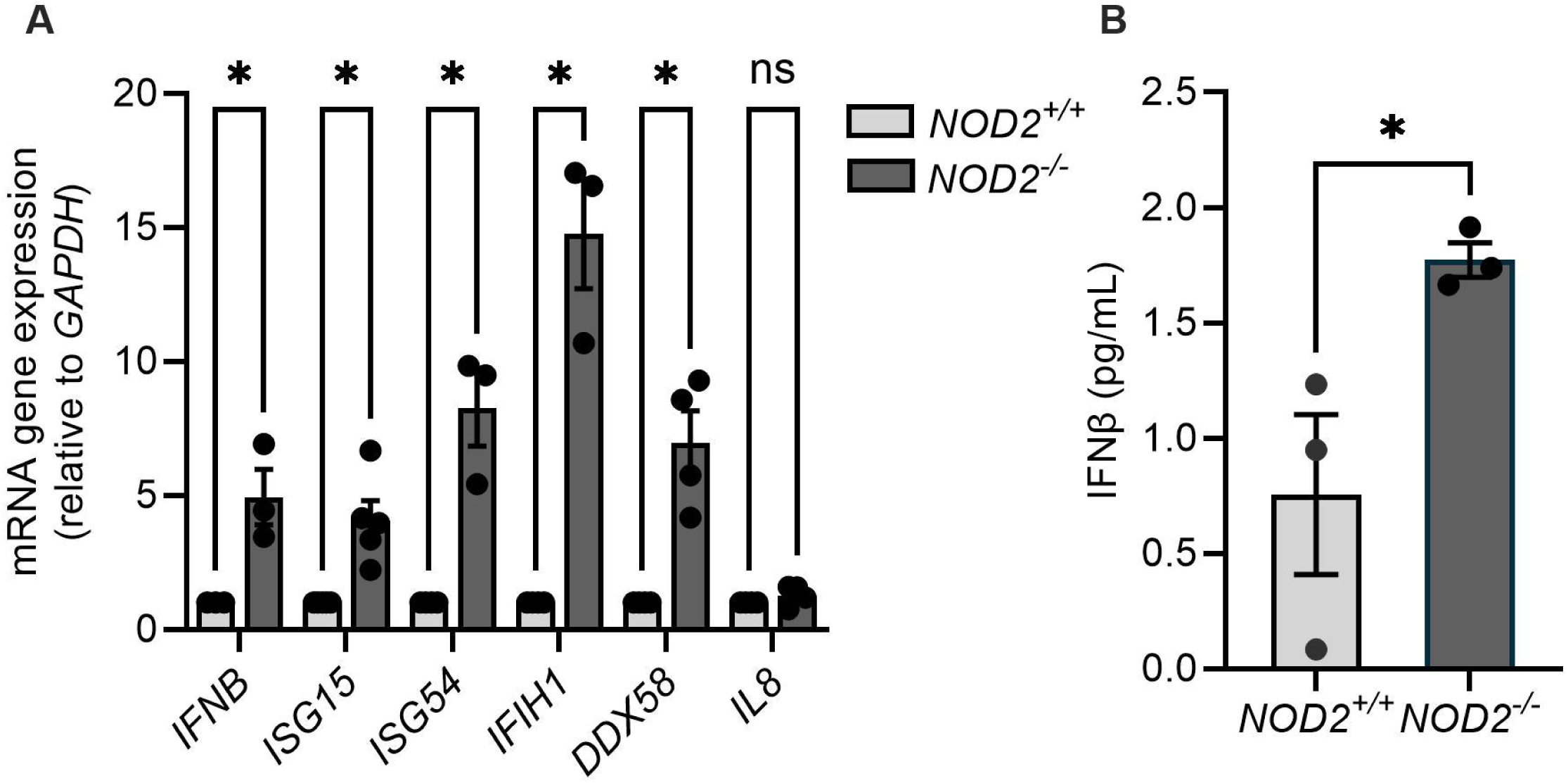
NOD2 suppresses type I interferon expression. *NOD2^+/+^* or *NOD2^−/−^*HCT116 cells were plated and RNA and supernatant collected 24 hours later. (A) *IFNB*, *ISG15, ISG54*, *IFIH1*, *DDX58*, and *IL8* gene expression was examined by qRT-PCR (N=3-5, duplicates). \**p*<0.05, multiple unpaired *t*-test. (B) IFNβ production was measured by ELISA (N=3, duplicates). \**p*<0.05, unpaired *t*-test.

### Silencing of MDA5 restores CVB3 replication in *NOD2^−/−^* cells

We hypothesized that increased expression of MDA5 and/or RIG-I in *NOD2^−/−^* cells would inhibit viral replication since MDA5 and RIG-I are viral PRRs that activate T1IFN ^27, 51^. MDA5 is especially critical for initiating T1IFN signaling during CVB3 replication ^26–29^, and the effects of RIG-I on CVB3 infection are not fully defined, but could play a role ^40^. To determine whether reducing *IFIH1* or *DDX58* expression would enhance CVB3 replication in *NOD2^−/−^* cells, *NOD2^+/+^* and *NOD2^−/−^* cells were transfected with control, *IFIH1* or *DDX58* siRNAs and then infected with CVB3 for 24 hours. Depletion of *IFIH1* but not *DDX58* significantly increased CVB3 replication in *NOD2^−/−^* cells (Fig. 5A and Fig. S4A), suggesting that negative regulation of MDA5 expression by NOD2 promotes CVB3 replication. We did not observe a full rescue of CVB3 replication by *IFIH1* knockdown, which could suggest that remaining levels still inhibited CVB3. Indeed, *IFIH1* and *DDX58* were well depleted by the siRNAs, but some remaining protein was detectable in *NOD2^−/−^* cells treated with either siRNA (Fig. S4B). Cell viability was not affected by siRNA (Fig. S4C). To determine whether depletion of MDA5 would also result in reduced expression of the T1IFN response, we measured mRNA gene expression of *IFNB*, *ISG15*, *ISG54* and *IFIH1* and *IL8* (Fig 5B). All genes were significantly decreased, except *IL8*, indicating that MDA5 controls the T1IFN response in HCT116 cells.

**Figure 5.**
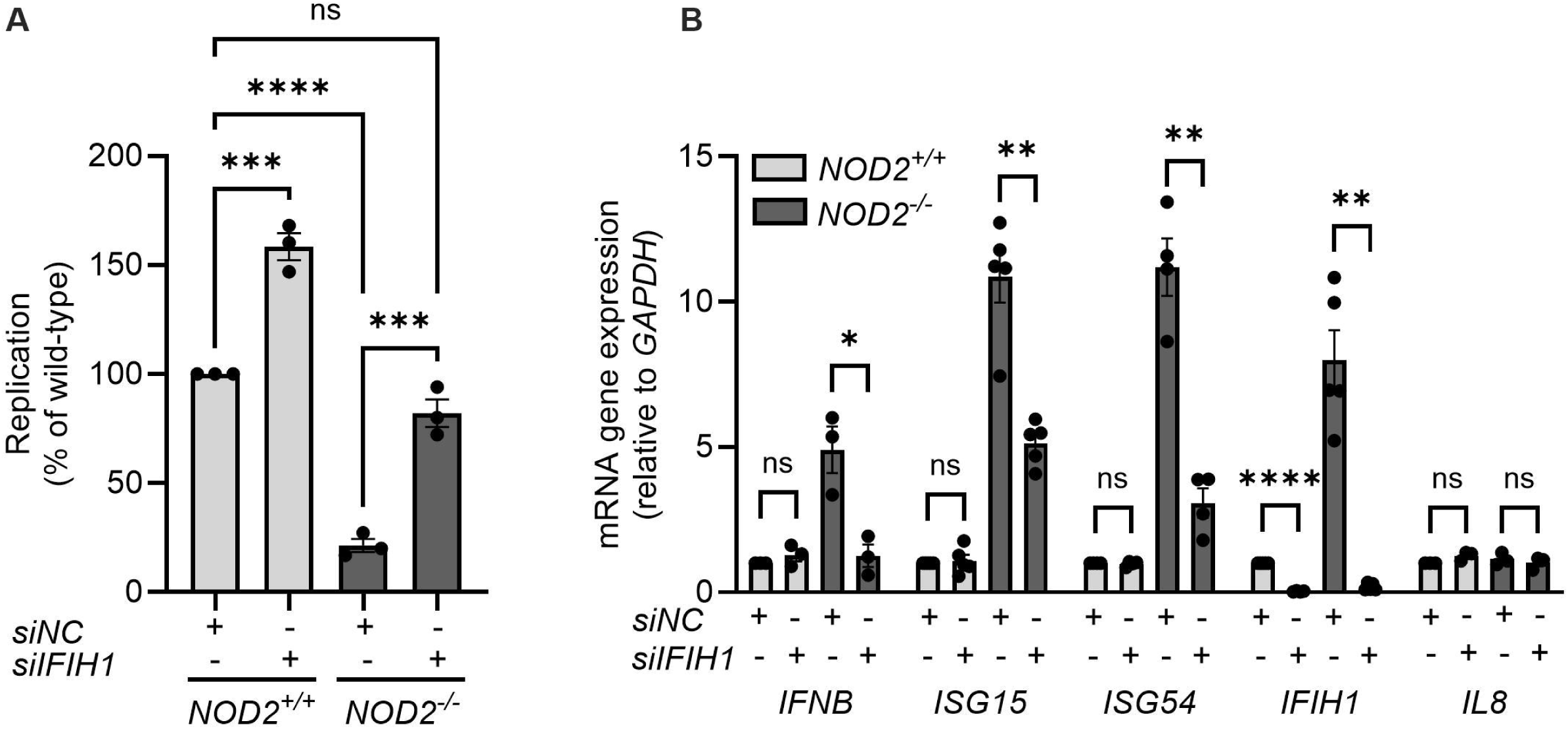
MDA5 regulates CVB3 replication in *NOD2^−/−^* HCT116 cells. *IFIH1* was siRNA depleted (10 nM) in *NOD2^+/+^* and *NOD2^−/−^* HCT116 cells for 2 days. (A) Cells were infected with CVB3 (MOI=0.5) for 24 hours and virus was quantified by plaque assay (N=3, duplicates). \*\*\**p*<0.001, \*\*\*\**p*<0.0001, ns=not significant, two-way ANOVA. (B) RNA was collected from uninfected cells and quantified for *IFNB*, *ISG15, ISG54*, *IFIH1*, *DDX58*, and *IL8* gene expression by qRT-PCR (N=3-5, duplicates). \**p*<0.05, \*\**p*<0.01, \*\*\*\**p*<0.0001, ns=not significant, multiple unpaired *t*-test.

### Stimulation with the MDA5 ligand PolyI:C increases the T1IFN response in *NOD2^−/−^* cells

To directly investigate whether increased MDA5 expression results in an increased T1IFN response when stimulated, we transfected HCT116 cells with the MDA5 ligand polyinosinic:polycytidylic acid (polyI:C). Unlike naked polyI:C, which mainly activates Toll-like receptor 3 (TLR3), transfected polyI:C can stimulate MDA5-dependent production of T1IFNs ^52^. Compared with the *NOD2^+/+^* IECs, NOD2 deficiency led to a stronger upregulation in the expression of ISGs in response to polyI:C stimulation (Fig. 6A). Similarly, supernatant collected from polyI:C stimulated *NOD2^−/−^* IECs contained higher levels of IFNβ (Fig. 6B), suggesting that NOD2 can suppress T1IFN production (Fig. 6).

**Figure 6.**
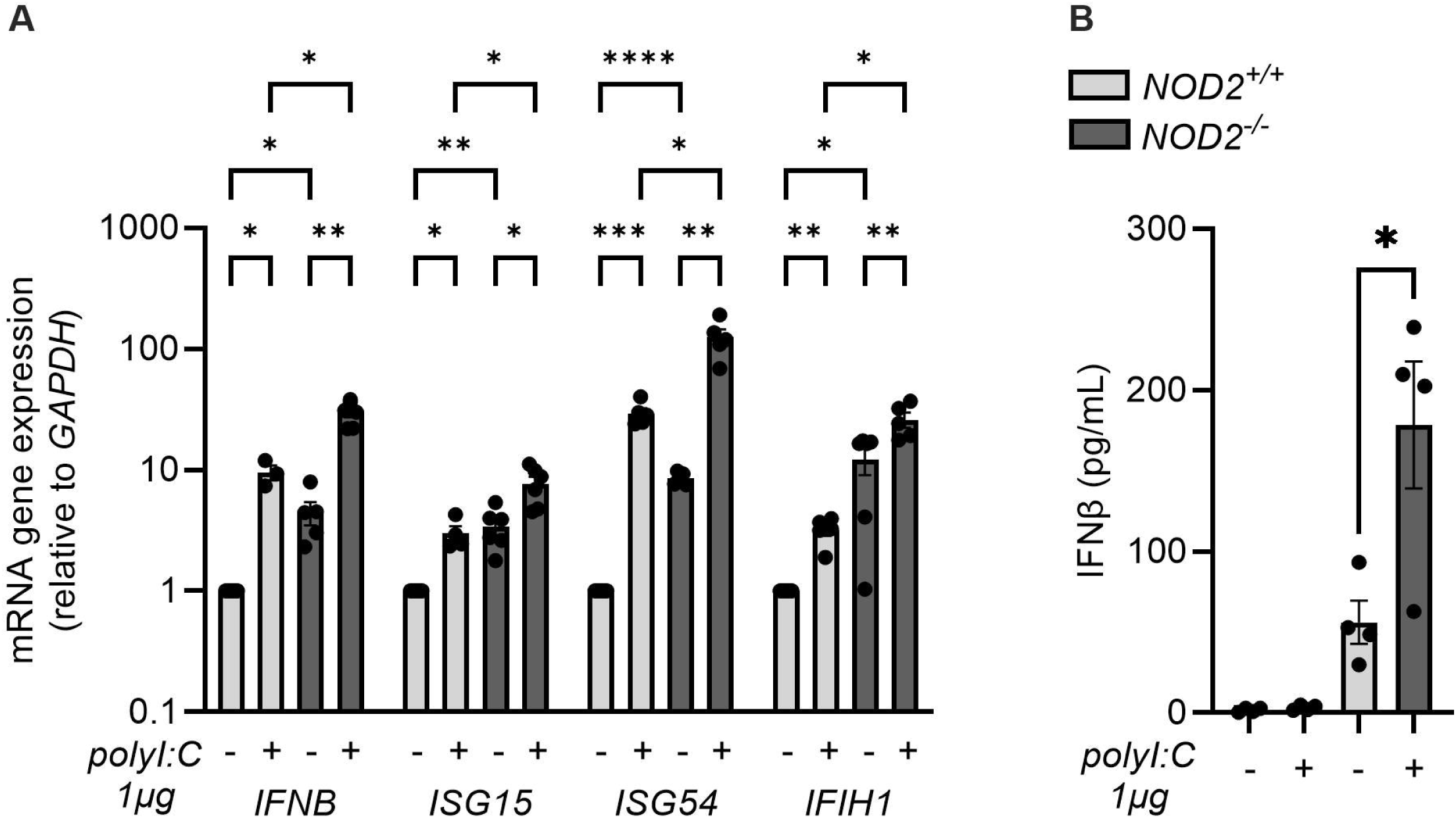
Stimulation with the MDA5 ligand PolyI:C increases the T1IFN response in *NOD2^−/−^* cells. *NOD2^+/+^* or *NOD2^−/−^* HCT116 cells were transfected with 1 µg/mL of polyI:C with PEI for 3 hours (A) and *IFNB*, *ISG15, ISG54*, and *IFIH1* gene expression was examined by qRT-PCR (N=3-5, duplicates). \**p*<0.05, \*\**p*<0.01, \*\*\**p*<0.001, \*\*\*\**p*<0.0001, two-way ANOVA, or for 24 h (B) IFNβ production was measured by ELISA (N=4, duplicates). \**p*<0.05, paired *t*-test.

### CVB3 replication depends on NOD2-mediated mitophagy

Viral RNA recognition by MDA5 leads to MDA5 interaction with MAVS ^53^. MDA5 binding to MAVS results in the formation of MAVS protein aggregates on the surface of mitochondria, which is necessary to attract other signaling molecules, including TBK1 and IRF3, to form the MAVS signaling complex and subsequent induction of IFNα and β expression ^53^. IFNα/β bind to the αβ receptor (IFNAR) to activate JAK/STAT signaling, which induces expression of ISGs to establish an antiviral cellular state ^24^. To prevent the detrimental effects of excessive innate immune signaling, cells degrade the MAVS signaling complex via mitophagy ^48^. NOD2 plays important roles in autophagic responses against invading bacteria, and *Ripk2^−/−^* cells and mice have increased reactive oxygen species (ROS) production indicative of accumulation of damaged mitochondria and defective mitophagy ^10, 54, 55^. To investigate whether NOD2 in IECs contributes to mitophagy, and consequently inhibits MDA5-mediated T1IFN responses, we initially measured mitochondrial superoxide production in *NOD2^+/+^* and *NOD2^−/−^* cells. At baseline levels, *NOD2^−/−^* cells produced significantly more superoxide compared to *NOD2^+/+^* cells, suggestive of dysfunctional mitochondria in *NOD2^−/−^* cells (Fig. 7A). To test whether mitophagy interferes with expression of *IFNB* and ISGs, we treated *NOD2^+/+^* and *NOD2^−/−^* cells with the mitophagy inducer carbonyl cyanide *m*-chlorophenyl hydrazone (CCCP) and performed qRT-PCR (Fig. 7B). The T1IFN response is significantly reduced in *NOD2^+/+^* cells treated with CCCP, while, as hypothesized, induction of mitophagy with CCCP in the *NOD2^−/−^* cells had no significant effect on the T1IFN response, suggesting that the MAVS/MDA5 signaling complex is degraded via NOD2-dependent mitophagy resulting in suppression of the T1IFN response. Finally, induction of mitophagy with CCCP increased viral replication only in *NOD2^+/+^* cells and not in the *NOD2^−/−^* cells, indicating that NOD2 promotes mitophagy and dampening of the antiviral T1IFN response (Fig. 7C). Overall, our data demonstrate that NOD2 plays a role in suppressing MDA5-mediated T1IFN responses, likely through mitophagy regulation, which in turn makes IECs more susceptible to CVB3 replication (Fig. 8).

**Figure 7.**
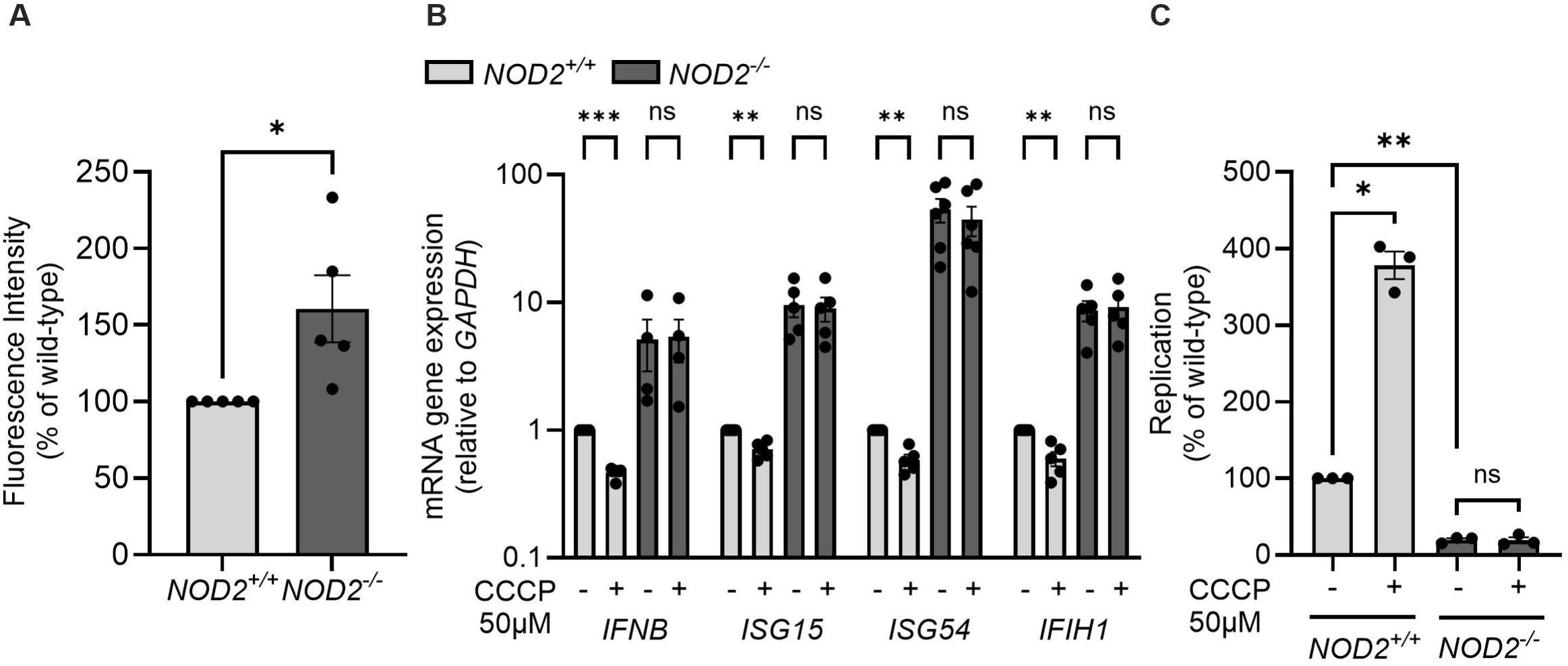
CVB3 replication depends on NOD2-mediated mitophagy. (A) *NOD2^+/+^* or *NOD2^−/−^*HCT116 cells were stained with 1 µM Mitosox Green reagent and fluorescence was measured as an indicator of mitochondrial superoxide production. (N=5, duplicates). \**p*<0.05, paired *t*-test. (B) *NOD2^+/+^* or *NOD2^−/−^* HCT116 cells were stimulated with 50 µM of CCCP and RNA was harvested after 2 hours, and *IFNB*, *ISG15, ISG54*, and *IFIH1* gene expression was examined by qRT-PCR (N=4-6, duplicates). \*\**p*<0.01, \*\*\**p*<0.001, ns=not significant, multiple paired *t*-test. (C) *NOD2^+/+^* or *NOD2^−/−^* HCT116 cells were pretreated for 20 minutes prior to CVB3 infection (24 hours) at an MOI=0.5. Virus was quantified by plaque assay and displayed as percent virus compared to wild-type control (N=3, duplicates). \**p*<0.05, \*\**p*<0.01, ns=not significant, one-way ANOVA.

**Figure 8.**
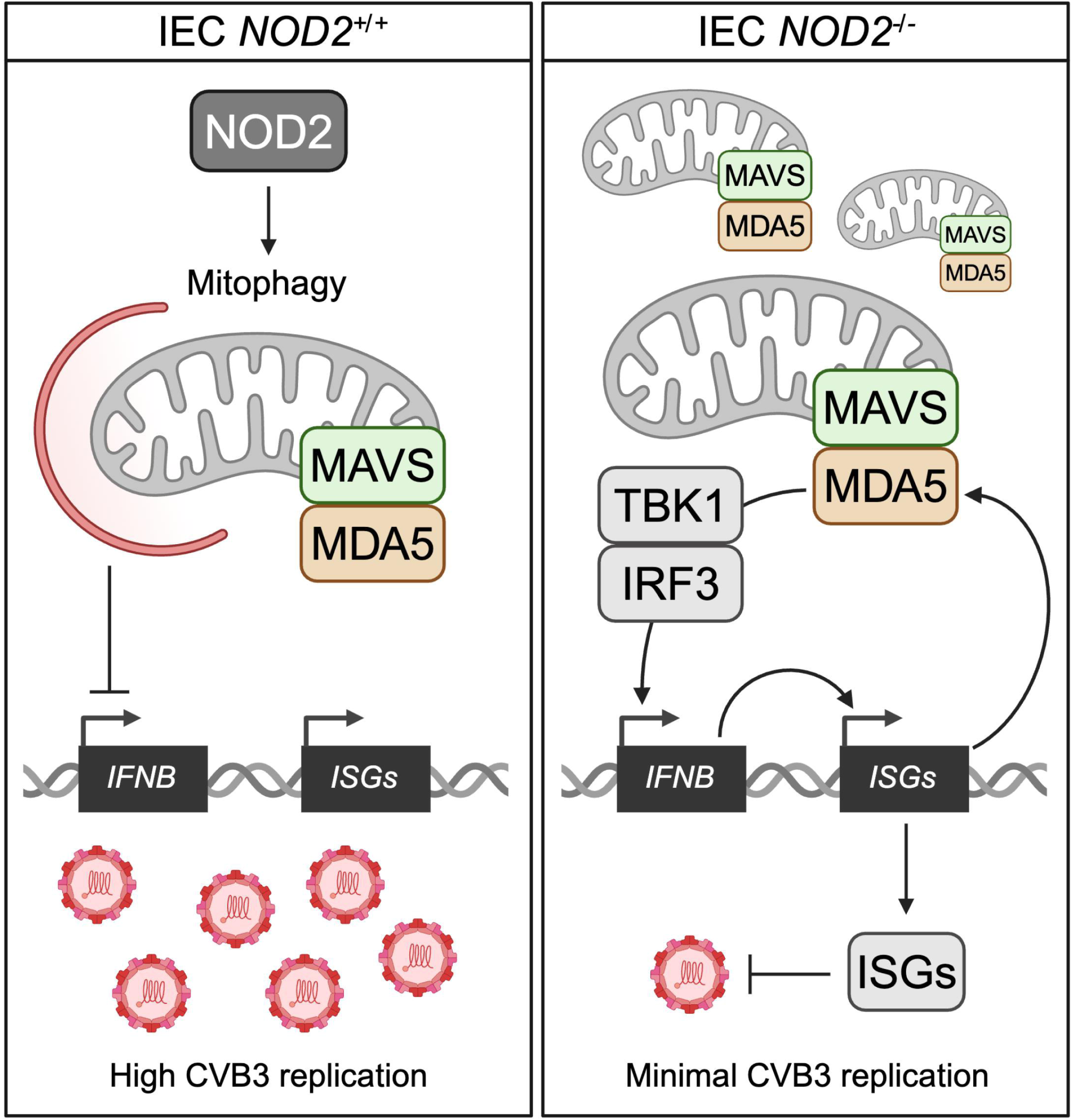
Model summarizing the main findings of this work. NOD2 promotes CVB3 replication by suppressing T1IFN expression and MDA5 activation through NOD2-mediated mitophagy. *NOD2^−/−^* IECs have increased expression of *IFNB* and ISGs, including MDA5, and reducing MDA5 expression in *NOD2^−/−^* IECs rescues CVB3 replication. In *NOD2^+/+^*IECs, T1IFN responses are decreased and CVB3 replication is increased when mitophagy is induced. However, mitophagy induction in *NOD2^−/−^* IECs does not affect the T1IFN response or CVB3 replication, indicating a role for NOD2 in mitophagy.

## Discussion

Rapid immune responses to pathogens are crucial to control and clear pathogenic infections. PRRs, like NOD-like receptors (NLRs), RLRs, TLRs and C-type lectin receptors (CLRs) are the first lines of defense against invading pathogens ^56^. However, hyperactivation of some of these molecules can lead to autoimmunity, and therefore, careful regulation of PRRs is critical to control inflammation ^57^. Indeed, polymorphisms in the PRR NOD2 are the most common mutations associated with autoimmune inflammatory bowel diseases ^58–60^. Furthermore, unchecked TLR2-induced inflammation predisposes mice to colitis, and NOD2 suppresses TLR signaling in an IRF4-dependent manner ^61–66^. Thus, although NOD2 elicits pro-inflammatory responses following stimulation by many different ligands, it also downregulates strong NFκB signaling induced by TLRs. Counter to this, NOD2 can also work synergistically with TLRs ^57^ and other innate immune pathways, such as T1IFN ^67^. NOD2 enhances T1IFN responses during some viral and bacterial infections ^67^. However, recent evidence demonstrated that NOD2 also suppressed T1IFN responses, which promoted infection with diverse viruses ^15^. In line with these latter findings, we demonstrate that NOD2 controls inflammation via mitophagy to reduce expression of MDA5 and other ISGs, which in turn facilitated CVB3 replication. Poliovirus replication was also enhanced by NOD2, likely via the same mechanism. NOD2 plays a crucial role not only in immune signaling, but also in maintaining intestinal homeostasis ^68^. Consequently, our findings indicate that CVB3 replication is enhanced due to NOD2-mediated suppression of MDA5, allowing CVB3 to escape T1IFN innate immune responses.

Our data indicate that the reduced viral load in *NOD2^−/−^*IECs is a result of CAR-independent uptake and is mediated by the MDA5-dependent T1IFN response. A previous report, however, demonstrated that CAR expression is reduced in the absence of NOD2, and suggested that CAR-dependent viral uptake is decreased resulting in the reduced viral load in *Nod2^−/−^* mice ^16^. CAR expression was measured in the myocardium of *Nod2^−/−^* CVB3 infected mice and in the murine cardiomyocyte cell line HL-1. Possibly, CAR expression in IECs is differentially regulated and NOD2-mediated downregulation of CAR is cell-type/tissue specific. CAR is ubiquitously expressed in epithelial tight junctions ^69^, and it is protective during intestinal inflammation by maintaining tight junction protein components and structure ^70^. Since it has been demonstrated that CAR is negatively regulated by inflammation ^70^, it is possible that in the absence of NOD2-mediated inflammation, CAR expression is not altered in IECs. CAR-dependent maintenance of the intestinal barrier function is critical to the host, and thus CAR expression may be more tightly regulated in the intestine (than the heart), which could prevent members of the resident microbiota from penetrating the intestinal epithelial barrier and disseminating to distant organs.

NOD2 and MDA5 both contain CARD domains, and they may interact with each other, thereby blocking MDA5-MAVS interactions that would normally stimulate T1IFN gene expression and further increase *IFIH1* levels. NOD2-MDA5 interactions might preclude oligomerization of MDA5, which is critical for its activation ^27^. In fact, NOD2 and RIG-I directly interact via the RIG-I CARD domains and multiple regions of NOD2 and exhibit regulatory crosstalk ^71^. Morosky *et al.* demonstrated that NOD2 negatively regulates RIG-I induction of T1IFN, and RIG-I reduces NOD2 activation of NFκB ^71^. Our data show similar negative regulation of RIG-I expression by NOD2, however silencing of RIG-I did not rescue CVB3 replication in *NOD2^−/−^* IECs, indicating RIG-I does not contribute to CVB3-induced innate immune responses in these cells. The increased RIG-I expression in the absence of NOD2 may be a consequence of the increased production of IFNβ, since *DDX58* is an interferon-stimulated gene ^72^, and not necessarily because of a direct inhibitory effect of NOD2 on RIG-I. It is conceivable that NOD2 and MDA5 interact similarly, which could lead to decreased MDA5 activation and T1IFN.

Our study, along with Limonta *et al*., demonstrate that NOD2 can promote viral infections by suppressing ISGs. This is shown for CVB3 and poliovirus, ZIKV, MAYV, DENV, SARS-CoV-2 and CVB5 ^15^. Conversely, prior studies demonstrated that NOD2 can inhibit viral infections by facilitating T1IFN activation during infections with human and mouse CMV ^13, 14^, RSV and VSV ^9^, IAV ^9–11^ and FMDV ^12^. How is NOD2 playing such divergent roles with different viruses? Viruses that utilize NOD2 for replication are all positive-sense RNA viruses, whereas those inhibited by NOD2 are mostly negative-sense RNA viruses, with the exception of the DNA virus CMV and positive-sense virus FMDV. Both positive and negative strand RNA viruses produce a dsRNA replicative intermediate, but interestingly, dsRNA is abundantly produced during positive-strand virus infections, and not during negative-strand virus infections ^73^ indicating that MDA5 might be more readily activated by positive strand RNA viruses. We and others ^15^ observed NOD2 inhibition of T1IFN in response to positive strand RNA viruses, but NOD2 facilitated T1IFN responses to negative strand viruses and ssRNA ^9^. Whether NOD2 responds differently to the presence of dsRNA versus ssRNA in the cytoplasm is unknown, but it could help explain its discrepant functions. Cell type specific functions of NOD2 likely do not explain this paradox as Limonta *et al.* demonstrated NOD2 reduction of T1IFN in IFNα-stimulated A549 epithelial cells treated +/-NOD2 signaling inhibitors ^15^, whereas Sabbah *et al.* observed NOD2 promotion of T1IFN in RSV-infected A549 epithelial cells ^9^. Thus, the same cell type yielded vastly different effects of NOD2 during differing viral infections. Interestingly, other NLRs restrict RLR signaling, including NLRP12 ^74^ and NLRX1 ^75, 76^. NLRP12 interferes with TRIM25-mediated activation of RIG-I and promotes RIG-I degradation ^74^. NLRX1 inhibits T1IFN induction by RIG-I and MDA5 by blocking their interactions with MAVS ^75, 76^, which promotes viral replication; however, Moore *et al.* observed that NOD2 did not display the same inhibitory effect on *IFNB* gene expression as NLRX1 ^76^. Therefore, NLRs may represent a class of checkpoint inhibitors for RLRs to restrain excessive inflammation.

The direct mechanism underlying NOD2 suppression of MDA5 is unclear, but our data suggest that the downstream adapter RIPK2 is also involved since inhibition of RIPK2 with GSK583 reduced CVB3 replication to similar levels as knocking out *NOD2*. Our data further suggest that NOD2 is required for mitophagy and the control of mitochondrial superoxide production. Supporting our findings, activation of NOD2 promoted intestinal stem cell survival by inducing mitophagy and protecting against ROS-mediated cytotoxicity ^77^. Lupfer *et al*. demonstrated that RIPK2 phosphorylates the mitophagy inducer ULK1, resulting in decreased mitochondrial ROS production ^10^. The precise mechanism by which NOD2 contributes to mitophagy is still unknown and future research is needed to uncover the specifics. NOD2 plays a vital role in autophagy by recruiting the autophagy proteins ATG5, ATG7 and ATG16L1 to the autophagosome ^54, 55, 78^. It is possible that these proteins are also recruited to the mitophagosome in a NOD2/RIPK2-dependent manner, facilitating the formation of the mitophagosome and the subsequent degradation of damaged mitochondria along with the associated MDA5/MAVS signaling complex. Interestingly, CVB3 actively induces mitophagy to enhance viral replication potentially benefiting from NOD2-mediated mitophagy ^49^.

The increased baseline expression of T1IFNs and ISGs in *NOD2^−/−^*cells, even without MDP stimulation, can potentially be attributed to the release of mt-dsRNA from damaged mitochondria, which can activate MDA5 ^79^. Since NOD2 has a regulatory role in the intestine and can control mitophagy at homeostasis to prevent accumulation of damaged mitochondria, it is possible that in the absence of NOD2 the damaged mitochondria release mt-dsRNA in the cytosol to activate MDA5. Future studies will determine whether the cytosol of *NOD2^−/−^* IECs have higher mt-dsRNA levels than *NOD2^+/+^* IECs. MDA5 is implicated in several autoimmune diseases, including type 1 diabetes, Aicardi-Goutiéres syndrome (AGS) and systemic lupus erythematosus (SLE). Single nucleotide polymorphisms in the MDA5 gene *IFIH1* likely stimulate chronic T1IFN that leads to the development of autoimmune interferonopathies ^27^. Furthermore, repetitive cellular RNA elements were shown to bind and stimulate MDA5, and therefore, cellular RNAs can trigger MDA5 activation ^79–83^. The MDA5 etiological link to autoimmunity underscores the importance of regulating MDA5 expression. Therefore, NLRs such as NOD2 may play a role in managing pathogenesis induced by microbes and hyper-inflammatory states caused by overactive MDA5.

Altogether, our data demonstrate that NOD2-driven inhibition of MDA5 suppresses ISG expression, which allows more efficient replication of enteroviruses CVB3 and poliovirus. Since CVB3 and poliovirus are enteric viruses that are thought to initiate infections in IECs, inhibition of MDA5 by NOD2 in IECs could facilitate their replication within the intestine and lead to enhanced dissemination and pathogenesis. Future studies to address this would be of interest although oral infection mouse models are limited for CVB3 and poliovirus, and they usually require *IFNAR1* deficiency ^45, 84–86^, which precludes studying the effects of intestinal NOD2 on T1IFN during enterovirus infections. Whether NOD2 negatively regulates MDA5 in other tissues is unknown, but our data show that NOD2 also promotes CVB3 replication in MEFs, indicating that this phenomenon could be broadly applicable. Further understanding the mechanism how NOD2 modulates mitophagy and suppresses T1IFN responses may reveal new insights into how NLRs regulate RLRs under differing pathogenic conditions.

## Materials and Methods

### Cells

HeLa cells (a gift from Julie Pfeiffer, UT Southwestern Medical Center), HCT116 cells (a gift from Andrew Thorburn, UC Denver Anschutz Medical Campus) and HEK293T cells (a gift from Dohun Pyeon, Michigan State University) were cultured in DMEM high glucose (Cytiva or Gibco) + 10% FBS (Millipore Sigma) + L-glutamine (Gibco) at 37°C with 5% CO_2_. Caco-2 cells (a gift from Sean Colgan, UC Denver Anschutz Medical Campus) were cultured in IMDM high glucose (Cytiva) + 10% FBS (Millipore Sigma) + L-glutamine (Gibco) at 37°C with 5% CO_2_. MODE-K cells were cultured in DMEM high glucose (Cytiva or Gibco) + 10% FBS (Millipore Sigma) + L-glutamine (Gibco) + non-essential amino acids (Gibco) + sodium pyruvate (Gibco) at 37°C with 5% CO_2_. Mouse embryonic fibroblasts (MEFs) were generated from gravid *Nod2^+/+^* and *Nod2^−/−^* mice as described previously ^87^ and cultured in DMEM high glucose (Cytiva or Gibco) + 10% FBS (Millipore Sigma) + L-glutamine (Gibco) + antibiotic-antimycotic (Gibco) at 37°C with 5% CO_2_.

### Generation of *NOD2^−/−^* HCT116 cells

No template control (NTC, wild-type) or NOD2 guide RNAs (gRNA) were cloned into lentiCRISPRv2-puro vector by the Functional Genomics Facility (UC Denver Anschutz Medical Campus; NOD2 guide RNA: TGCCACATGCAAGAAGTATATGG [Sigma CRISPR clone reference]). To generate lentiviruses expressing either NTC or NOD2 gRNAs, 3 μg lentiCRISPRv2-puro-NTC or lentriCRISPRv2-puro-NOD2 vectors were co-transfected with 750 ng pVSV-G and 2.25 μg pRΔ8.2 (gifts from Dohun Pyeon, Michigan State University) per 10 cm dish of 293T cells using Lipofectamine 2000 (30 μL per plate) in a total volume of 750 μL/plate in Optimem (Gibco) following Lipofectamine 2000 manufacturer’s protocol. Media (DMEM + 10% FBS) was changed the following day and cells incubated at 37°C. At 48 and 72 hours, media was harvested and cell debris was removed by centrifugation at 500 x g for 5 minutes. Supernatants containing lentiviruses were aliquoted and frozen at -80°C.

HCT116 cells were plated at 2 × 10^5^ cells/well in a 24 well plate to reach confluency of 70% the following day. Cells were infected with 300 μL NTC or NOD2 lentiviruses + polybrene (8 μg/mL) + 200 μL of media and incubated at 37°C for 24 hours. The next day, cells were confluent and split into a 6 well plate. Four days post-infection, media was replaced with 4 μg/mL puromycin media to select for CAS9/gRNA-expressing cells. Puromycin media was maintained for two weeks, and cells were plated at a low confluency to pick clones. Clones were picked by disc trypsinization of individual colonies that were transferred to 96 well plates and expanded. Depletion of NOD2 was functionally assessed by 1 or 10 μg/mL muramyl dipeptide (MDP, Invivogen) stimulation of cells. One clone lacked MDP stimulation of *IL8* gene expression indicating that NOD2 was non-functional. This HCT116 clone is referred to as “*NOD2^−/−^*” while the NTC HCT116 cells are referred to as “*NOD2^+/+^*” (or wild-type). Cell growth of each clone was examined using CellTiterGlo (Promega) following manufacturer’s protocol.

To confirm CAS9-mediated mutations in *NOD2^−/−^* cells, DNA was purified, amplified, and subjected to sequencing following the Takara Bio USA Guide-It Indel Identification Kit protocol. *NOD2^+/+^* and *NOD2^−/−^* HCT116 cellular DNA was amplified following the manufacturer’s protocol using primers specific to the region aligning with the NOD2 guide RNA (pUC19-NOD2-F: CGGTACCCGGGGATCACTGTCCTCAAAGTGCACAGCTTGT; pUC19-NOD2-R: CGACTCTAGAGGATCCAGGGTGGTCAAGGAGTAACTGGAA) using *Homo sapiens* NOD2 gene NC_000016 as a reference. Amplicons were gel purified (Nucleospin gel and PCR Clean Up) and cloned into pUC19 using the manufacturer’s provided In-fusion cloning master mix. 2.5 μL of the resulting products were transformed into 50 μL of Stellar Competent Cells (Clontech) following manufacturer’s protocol and plated on LB Agar plates with 100 μg/mL ampicillin. Colonies were screened by colony PCR using the provided Colony PCR Forward and Reverse Primers (Takara Bio USA) to confirm that the amplicons were incorporated into pUC19. Positive products were purified using GeneJet PCR Purification Kit (Thermo Scientific) and submitted for sequencing with the Colony PCR Forward Primer (Takara Bio USA) to Quintarabio (Denver, CO). Sequencing results were aligned to *Homo sapiens NOD2* transcript variant 1 (NM_022162) mRNA to identify the mutations within *NOD2* exons.

### Viruses, Infections and Plaque Assays

CVB3-H3 was generated from the pCVB3-H3 plasmid (a gift from Julie Pfeiffer, UT Southwestern Medical Center) by in vitro transcription (mMessage mMachine T7 Kit, Life Technologies). Poliovirus type 1 (Mahoney) was provided by David Barton (UC Denver Anschutz Medical Campus) and generated as previously described ^88^. 10 μg CVB3-H3 RNA was used to transfect a 10 cm plate of HeLa cells (∼80% confluency) using the Qiagen TransMessenger Transfection Reagent following manufacturer’s protocol. Cells were observed until significant cytopathic effect (CPE) was reached (∼36 hours), at which cells were harvested by trypsinization and centrifugation at 500 *x g* for 5 minutes. Cells were resuspended in PBS+ (PBS supplemented with 100 μg/ml MgCl_2_ and 100 μg/ml CaCl_2_) and freeze-thawed three times to lyse cells and release progeny virions. This virus stock was expanded to achieve a higher titer stock by infecting a 15 cm plate of HeLa cells at a high MOI and harvesting after one replication cycle (∼7 hours post-infection).

Cell plating for infections was done as follows for each cell line, 24 well: HCT116: 3.8 × 10^5^, MODE-K: 1 × 10^5^; 12 well: HCT116: 9 × 10^5^; 6 well: MEF: 1.5 × 10^5^. Cells were incubated at 37°C overnight prior to experimentation. CVB3-H3 was diluted in full media (see “Cells”) to achieve the indicated MOIs and used to infect cells for 30 minutes at 37°C. Infections volumes were 125 μL for 24 well plates and 150 μL for 12 well plates. After 30 minutes with intermittent rocking, infection mixes were removed, cells were washed twice with PBS+, fresh media was added and infections allowed to proceed at 37°C. Supernatants and/or cells were harvested for plaque assay quantification and stored at -80°C. For cell-associated virus, cells were harvested by trypsinization, diluted 1:1 in PBS+ and centrifuged at 4000 *x g* for 4 minutes. Resulting supernatants were removed, cell pellets were resuspended in 100 μL of PBS+ and freeze-thawed three times prior to quantification by plaque assay. For inhibitor treatments and mitophagy induction, cells were pre-treated for 20 minutes at 37°C with 0.1% DMSO (Fisher BioReagents) control or GSK583 (Cayman Chemicals), or 50 µM CCCP (Thermo Fisher Scientific). Infections were performed in the presence of DMSO or treatment for 30 minutes at 37°C, cells were washed twice with PBS+ and full media with DMSO or inhibitor was added for the experiment duration. For mitophagy induction, cells were infected without CCCP. Cell viability assays were performed by LDH release assay following manufacturer’s protocol (CytoTox96, Promega).

Plaque assays were performed on either cell supernatants or cell-associated virus. Dilutions were made in PBS+ and HeLa cells (2.7 × 10^5^) were infected with 200 μL of each dilution for 30 minutes at 37°C in 6 well plates (Costar). Infection mixes were removed and 2 mL of agar overlays were added per well. Agar overlays consisted of 2X DME, generated using DME powder (Gibco, manufacturer’s protocol amended to make 2X) with 10% FBS (Sigma) and 1X antibiotic-antimycotic (Gibco), mixed 1:1 with 2% Bacto agar (BD). Plaque assays were incubated for 2 days at 37°C after which agar overlays were removed and cells were stained with crystal violet staining solution (20% ethanol, 0.1% crystal violet). Virus replication displayed as PFU/mL is indicative of the amount of cell-associated virus/mL from the 100 μL aliquot or directly from cell supernatants. Virus replication displayed as a percent of wild-type is normalized to the average of the wild-type values multiplied by 100.

### Virus Binding Assays

*NOD2^+/+^* and *NOD2^−/−^* HCT116 cells were plated at 2 × 10^5^ cells/well in two separate 48 well plates and set at 37°C over-night. The following day, equivalent amounts of isotype IgG control (GTX35009, Genetex) or anti-mouse CAR (05-644, clone RmcB, Millipore) were added to cells at 1:125 for 40 minutes at 37°C. After the incubation, one plate was kept at 37°C and another plate was pre-chilled on ice for 10 minutes. Cells were then infected at MOI=0.5 in warm (37°C) or cold (6°C) full media in the presence of isotype control or CAR antibody and set at 37°C for 30 minutes. Infection mixes were removed, cells were washed twice with PBS+ and full media was added for the duration of the 24-hour infection. At 24 hpi, cells were collected for cell-associated virus and plaque assays as described above.

### NLR/RLR Agonist Stimulations

*NOD2^+/+^* and *NOD2^−/−^* HCT116 cells (9 × 10^5^/12 well) were stimulated with water control, 1 or 10 μg/mL MDP, or 1 µg/mL C12-iE-DAP (Invivogen). 1 µg/mL polyI:C (Invivogen) was transfected into *NOD2^+/+^* and *NOD2^−/−^* HCT116 cells using polyethylenimine (PEI) at a 4 to 1 ratio. Cells were washed with PBS and collected in TRIreagent (Molecular Research Center). RNA was extracted following manufacturer’s protocol (Molecular Research Center) and DNase treated following manufacturer’s protocol (TURBO DNase, Invitrogen) for 25 minutes at 37°C. 2.5 μL DNase Inactivation Buffer was added, incubated for 5 minutes at room temperature and centrifuged at 10,000 rpm for 2 minutes. RNA supernatants were quantified by spectrophotometry using the Nanodrop (Thermo Scientific).

### Reverse Transcription and Quantitative PCR

800 ng of RNA were used for reverse transcription to generate cDNA (Taqman RT, Applied Biosystems). For quantitative PCR, 2 μL of cDNA were used per well in a 10 μL reaction using SYBR Green PCR Master Mix (Applied Biosystems) and 0.2 μM each of forward and reverse primers (Table 1) in a 384-well plate (VWR) using the Applied Biosystems QuantStudio 7 Flex System. Fold change values were calculated using the -ΔΔCt method.

**Table 1.**
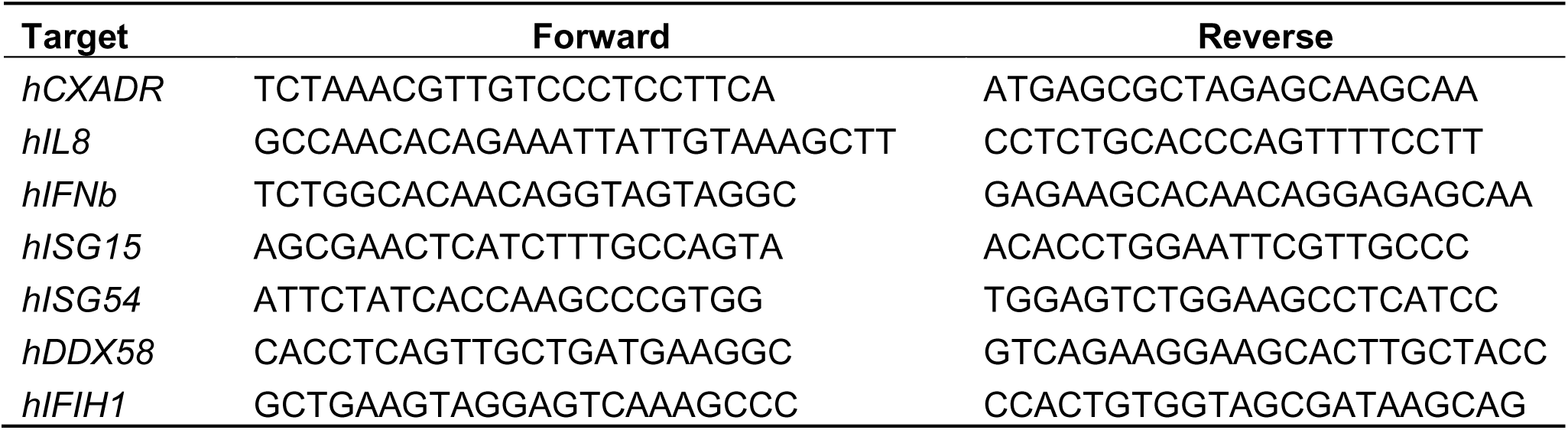
Primer sequences for qRT-PCR. (5’-3’)

### IFNβ ELISA

*NOD2^+/+^* and *NOD2^−/−^* HCT116 cells were plated and 24 hours later supernatant was collected to measure baseline IFNβ production. *NOD2^+/+^* and *NOD2^−/−^* HCT116 cells were transfected with 1 µg/mL polyI:C using PEI and 24 hours later supernatant was collected. Human IFNβ was measured using the DuoSet ELISA kit (R&D Systems, Cat# DY814-05) according to the manufacturer’s protocol.

### Superoxide production

*NOD2^+/+^* and *NOD2^−/−^* HCT116 cells were stained with 1 µM Mitosox Green reagent (Thermo Fisher Scientific) according to the manufacturer’s protocol. Bulk fluorescence was immediately read using the Tecan infinite M plex microplate reader at 488 nm.

### Immunoblots

Cells were collected in 2X Laemmli sample buffer (4% SDS (w/v), 20% glycerol, 120 mM Tris•HCl pH 6.8), boiled at 99°C for 10 minutes and stored at -20°C. Protein was quantified by BCA Protein Assay (Pierce) following manufacturer’s protocol for equivalent protein loading per sample (25 or 30 μg). Immunoblots were performed as previously described ^89^. Briefly, samples were diluted in 4X Laemmli sample buffer with 0.02% bromophenol blue dye, loaded on a 10% or 12% SDS-PAGE gel and ran in SDS-PAGE running buffer at 140V (Mini-PROTEAN Tetra Cell, Biorad). Transfers (Mini-Transblot, Biorad) were performed using methanol-activated PVDF membrane (Immobilon, Millipore). Membranes were blocked in 5% milk in TBST (137 mM sodium chloride, 20 mM Tris, 0.1% Tween-20) for 1 hour. Primary antibodies were diluted in 5% BSA (RPI) in TBST with 0.02% sodium azide (Sigma) and applied overnight at 4°C, rocking. Membranes were washed 3 x 5 minutes in TBST, and secondary antibodies (Jackson Immunoresearch) were added at 1:5000 in 5% milk/TBST for 1-2 hours at room temperature. Membranes were washed 3 x 5 minutes in TBST, developed using Clarity Western ECL (Biorad) or West Femto ECL (Pierce) and chemiluminescence was imaged by Syngene G:Box.

Antibodies used: mouse anti-CAR (sc-373791, clone E-1, Santa Cruz, 1:350); mouse anti-β-ACTIN (#3700, Cell Signaling Technologies, 1:5000); mouse anti-FLAG M2 (Sigma); rabbit anti-RIG-I (#3743, clone D14G6, Cell Signaling Technologies, 1:1000); rabbit anti-MDA5 (#5321, clone D74E4, Cell Signaling Technologies, 1:500).

### Immunofluorescent Staining and Microscopy

*NOD2^+/+^* and *NOD2^−/−^* HCT116 cells were plated at 1.9 × 10^5^ cells/well on Coverglass No. 1 coverslips (Fisherbrand) inserted into a 24 well plate (Greiner Bio-One) to achieve a confluency of ∼30%. 24 hours later, cells were washed and fixed in 4% paraformaldehyde (PFA) in PBS for 20 minutes at room temperature. PFA was removed and cells were washed 3 x 1 minute in PBS. To examine plasma membrane expressed CAR only, cells were not permeabilized prior to blocking in 10% normal goat serum (Thermo Scientific) for 1 hour. Rabbit anti-human CAR antibody (Cell Signaling, #16984) was added at 1:100 in 10% normal goat serum and rocked at 4°C over-night. Coverslips were washed 3 x 2 minutes in PBS, and goat anti-rabbit 488 secondary was added at 1:500 for 1 hour in 10% normal goat serum. Coverslips were washed 3 x 2 minutes in PBS, and Hoechst 33342 stain (Invitrogen) was added at 1:2000 in PBS for 5 minutes. Coverslips were washed 3 x 1 minute in PBS and mounted to slides (Fisherbrand) using Fluoromount-G (Electron Microscopy Sciences).

A Nikon TE2000-E Inverted Fluorescence Microscope or a Nikon Ti2 Epifluorescence Microscope with Hammamatsu BT Fusion Camera and Semrock Filters were used to image mounted coverslips. Image acquisition and analysis was performed using Slidebook 4.0 (Intelligent Imaging Innovations) or NIS-Elements software (Nikon). Z stack images (100X) were acquired for 488 and DAPI channels. Images were deconvolved, 488 intensity was quantified for whole images and determined per cell by dividing 488 intensity by the number of cells per image. Results are presented as percent CAR surface expression in which all values were normalized to the average of the wild-type values. Z-stacks from representative images were subjected to maximum image projection for publication.

## Statistical Analysis

Data analysis was performed with GraphPad PRISM. Data is shown as mean ± standard error of the mean (SEM). One-way ANOVAs and Student’s t-tests, were performed. More information about statistical analysis can be found in individual figure legends.

## Acknowledgements

We acknowledge Dr. Julie Pfeiffer (University of Texas Southwestern Medical Center) for providing the CVB3 plasmid, HeLa cells and protocols for generating CVB3. We are grateful to Drs. David Barton and Brian Kempf (University of Colorado Anschutz Medical Campus) for providing poliovirus and laboratory space to perform poliovirus experiments. Work in A.M.K-G.’s laboratory is supported by grants from the NIAID of the NIH under Award Numbers AI164154, AI173121 and AI188344.

