## Supplementary figures and images for "NOD2 modulates MDA5 signaling to promote coxsackievirus B3 replication"

### Figure S1

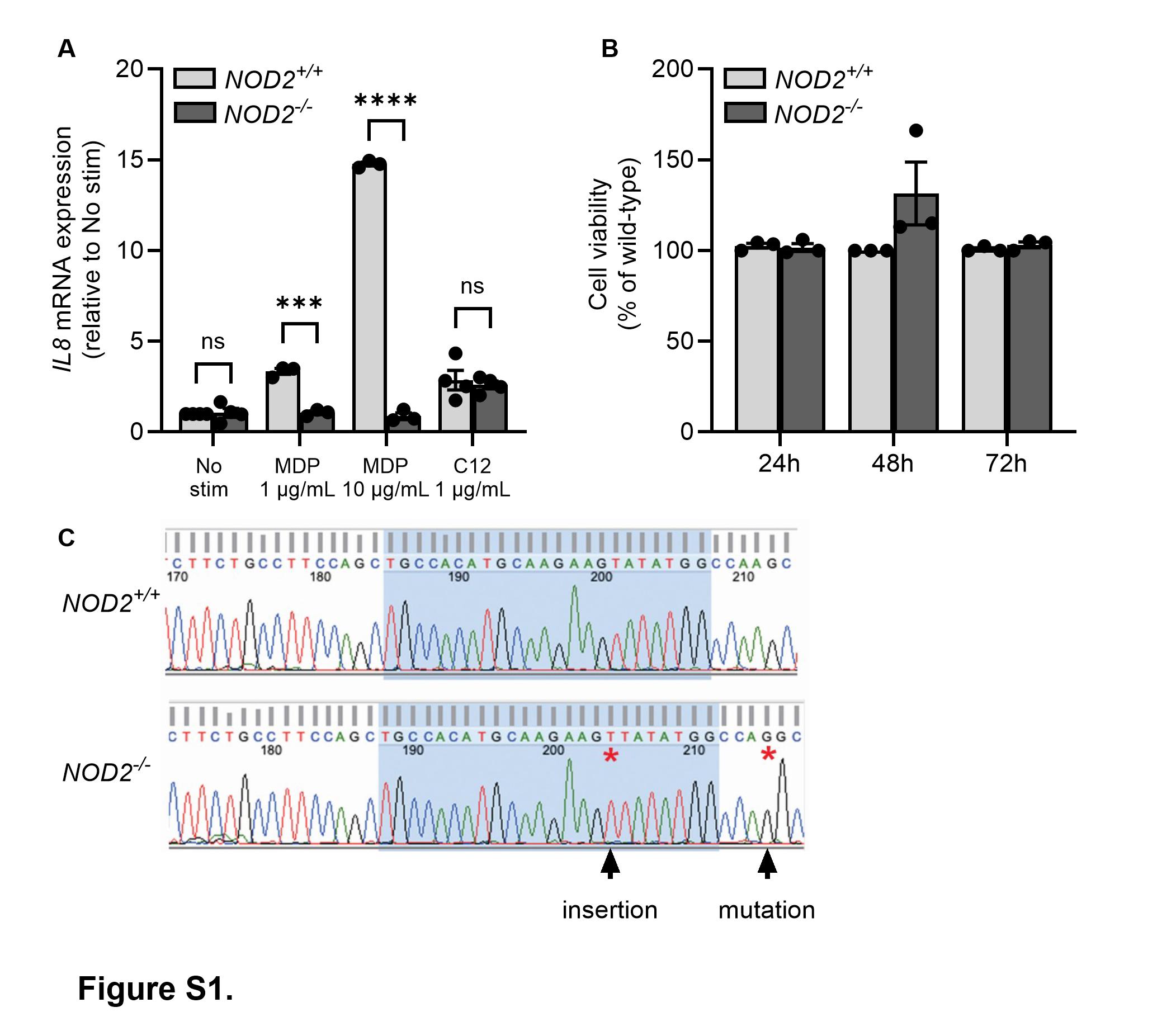

### Figure S2

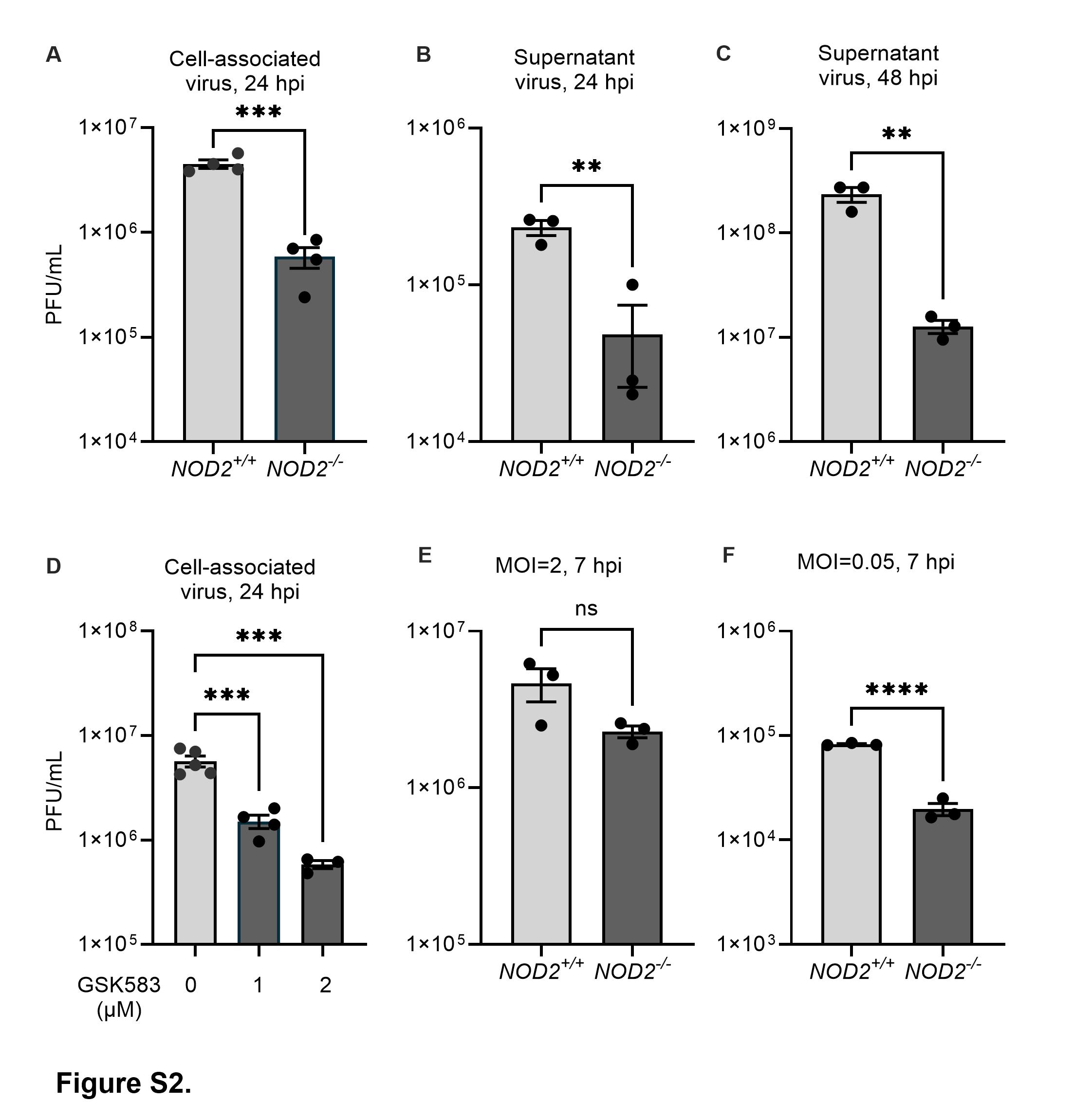

### Figure S3

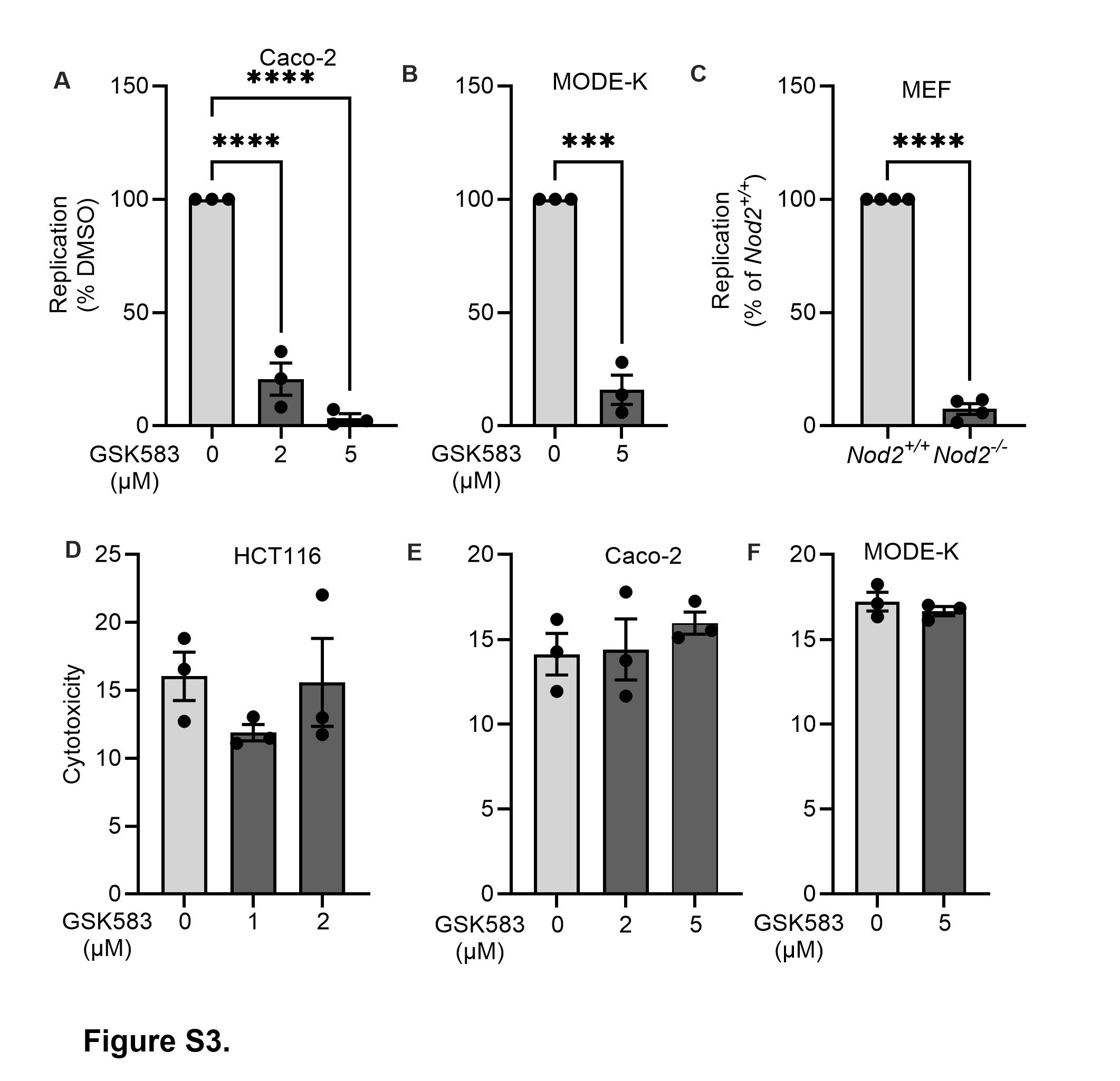

### Figure S4

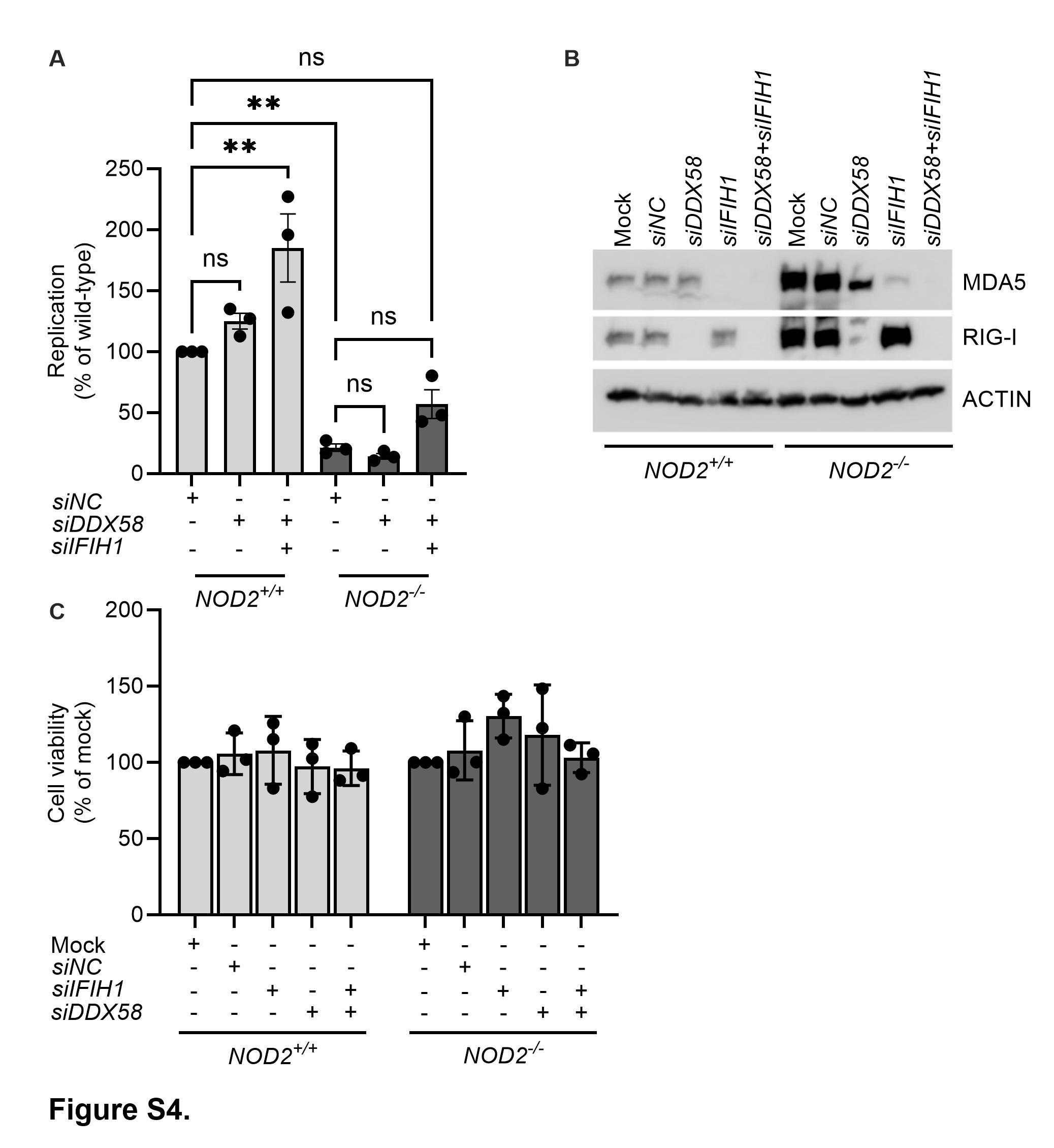
